# A Non-Coding Variant in *SLC15A4* Modulates Enhancer Activity and Lysosomal Deacidification Linked to Lupus Susceptibility

**DOI:** 10.1101/2023.07.28.551056

**Authors:** Manish Kumar Singh, Guru Prashad Maiti, HariKrishna Reddy-Rallabandi, Mehdi Fazel-Najafabadi, Loren L. Looger, Swapan K. Nath

**Affiliations:** Arthritis and Clinical Immunology Research Program, Oklahoma Medical Research Foundation, Oklahoma City OK, USA; Howard Hughes Medical Institute, Department of Neurosciences, University of California San Diego, La Jolla CA, USA

**Author notes:** Contributed equally (Manish Kumar Singh, Guru Prashad Maiti). **Correspondence**: Swapan K. Nath, Ph.D.

**Keywords:** SLC15A4, Luciferase, ChIP-qPCR, CRISPR, Lupus, Lysosomal acidification

## Abstract

Systemic lupus erythematosus (SLE) is a complex autoimmune disease with a strong genetic basis. Despite the identification of several single nucleotide polymorphisms (SNPs) near the *SLC15A4* gene that are significantly associated with SLE across multiple populations, specific causal SNP(s) and molecular mechanisms responsible for disease susceptibility are unknown. To address this gap, we employed bioinformatics, expression quantitative trait loci (eQTLs), and 3D chromatin interaction analysis to nominate a likely functional variant, rs35907548, in an active intronic enhancer of *SLC15A4*. Through luciferase reporter assays followed by chromatin immunoprecipitation (ChIP)-qPCR, we observed significant allele-specific enhancer effects of rs35907548 in diverse cell lines. The rs35907548 risk allele T is associated with increased regulatory activity and target gene expression, as shown by eQTLs and chromosome conformation capture (3C)-qPCR. The latter revealed long-range chromatin interactions between the rs35907548 enhancer and the promoters of *SLC15A4, GLTLD1*, and an uncharacterized lncRNA. The enhancer-promoter interactions and expression effects were validated by CRISPR/Cas9 knock-out (KO) of the locus in HL60 promyeloblast cells. KO cells also displayed dramatically dysregulated endolysosomal pH regulation. Together, our data show that the rs35907548 risk allele affects multiple aspects of cellular physiology and may directly contribute to SLE.

## 1 Introduction

Systemic lupus erythematosus (SLE) is a chronic autoimmune disease characterized by immune attack on the body’s tissues and organs, resulting in profound dyshomeostasis and damage to skin, joints, kidneys, cardiovascular system, and nervous system, among others. The disease is very heterogeneous, with each patient presenting uniquely, thus making diagnosis, treatment, and even basic understanding challenging [1]. Gender and ethnicity significantly influence the incidence and severity of SLE, with females and individuals of African, Hispanic, and Asian ancestry being both more prone to SLE and to severe manifestations like kidney disease and frequent hospitalization. Conversely, European-ancestry SLE patients tend to exhibit more skin manifestations but less renal involvement. Therefore, managing SLE and its comorbidities requires a comprehensive internal medicine approach and an understanding of its diverse underlying pathogenic mechanisms [2, 3, 4, 5].

SLE has a significant genetic component, evidenced by familial aggregation, sibling risk ratio, and twin studies [6]. SLE appears to be a highly polygenic disease, with >100 risk loci implicated to date [7, 8]. These SNPs and associated genes are involved in biological pathways related to tolerance, cell signaling, apoptosis, and other critical immune functions [9, 10]. Molecular pathway analysis has revealed different underpinnings of immune system homeostasis and SLE risk in diverse ethnic populations [9, 11, 12], emphasizing the importance of studying the genetics of complex disease in multiple ethnicities – both to elucidate fundamental biochemical pathways and to develop personalized diagnostics and treatments.

*SLC15A4* was discovered as an SLE risk locus in 2009 through GWAS on Chinese individuals. Since then, several SNPs have been associated with risk across Asian, European, Hispanic, and African ancestries [9]. However, despite strong genetic association, the actual functional SNP(s) and underlying biological mechanism(s) contributing to SLE pathogenesis are not understood [13, 14].

SLC15A4, also known as peptide/histidine transporter 1 (PHT1), is a member of the solute carrier family 15 of proton-coupled oligopeptide transporters [13, 14, 15, 16]. It is primarily located on the endolysosomal membrane of immune cells, where it transports histidine and bacterially-derived dipeptides such as the NOD2 ligand muramyl dipeptide (MDP). SLC15A4 is crucial for lysosomal acidification, as it generates a proton gradient through transport of the proton acceptor histidine. Moreover, SLC15A4 can recruit the adapter molecule “TLR adaptor interacting with SLC15A4 on the lysosome” (TASL), which regulates Toll-like receptor (TLR) function and promotes downstream signaling through type I interferon and Interferon Response Factor 5 (IRF5) [17]. Therefore, SLC15A4 plays a crucial role in regulating lysosomal function and innate immunity. SLC15A4 deficiency may promote lysosomal dysfunction and impaired autophagy, both associated with various autoimmune diseases, including SLE [18]. Consequently, these findings suggest that pharmacological intervention to restore or supplement the function of SLC15A4 may be a promising therapeutic approach for treating lupus and other endosomal TLR-dependent diseases [18, 19, 20, 21].

Here, we employ systematic bioinformatics to investigate SNPs in the *SLC15A4* locus, identifying rs35907548 as a likely regulatory variant. We show that this variant indeed underlies activity of a potent enhancer, chromatin interactions, expression of *SLC15A4* and nearby genes, and ultimately plays a decisive role in maintaining endolysosomal acidification, critical for proper function of immune cells. Our results finely localize SLE risk of the highly associated *SLC15A4* locus and give mechanistic insight into its function.

## 2 Materials and Methods

### 2.1 Bioinformatics

We utilized RegulomeDB [22] and Ldlink [23] to prioritize SNP regulatory potential. We added information on *cis*-regulatory elements, including chromatin accessibility (ATAC-seq and DNase-I hypersensitivity), canonical histone modifications (H3K27ac, H3K4me1, H3K4me3), and RNA polymerase II (Pol II) binding, from ENCODE [24] data from immune cell lines. Transcription factor binding sites were taken from ENCODE as well.

To evaluate cell type-specific expression quantitative trait loci (eQTLs) and target gene expression, we used data from ImmuNexUT (Immune Cell Gene Expression Atlas from the University of Tokyo) [25], encompassing 28 immune cell types from both healthy donors and individuals with 10 immune diseases. To broaden our analysis, we added other eQTL databases, including eQTLgen [26], which is tailored for European SNP-based eQTLs.

### 2.2 Cell lines

Dual-luciferase reporter assays were performed in HEK293, lymphoblastoid cell lines (LCLs; B-cell), Jurkat (T-cell), and HL-60 (promyeloblast). ChIP-qPCR assays were performed in LCLs (GM18624, GM18603). Chromatin-conformation capture (3C) experiments were in the same LCLs. CRISPR/Cas9 gene editing was in HL-60. Endolysosomal pH measurements were in wild-type and gene-edited HL-60. All cell lines were purchased from ATCC (American Type Culture Collection). All cells were tested for mycoplasma by PCR and used between passages 4-7.

### 2.3 Dual-luciferase assay

To investigate whether the rs35907548 region has enhancer activity, we utilized the well-established Dual-Luciferase Reporter Assay System. A detailed description of the method is provided elsewhere [27, 28]. Briefly, we cloned the 300 bp region, rs35907548 at the middle (chr12:129,282,013-129,282,312, hg 19) locus into pGL4.26 (Promega) and co-transfected with pGL4.74 (internal control) in HEK293, LCL, Jurkat, and HL-60 cells. After 24 hours, enhancer activity was measured using a Synergy H1 spectrophotometer (BioTek). Three experimental replicates were performed per cell type. Statistical significance was assessed by Student’s t-test using GraphPad PRISM; *p*-value < 0.05 was considered significant. We used two non-coding SNPs, rs12831705 and rs34616325, as negative controls for allele-specific luciferase activity.

### 2.4 Chromatin immunoprecipitation and quantitative PCR (ChIP-qPCR)

To investigate whether the rs35907548 region shows allele-specific binding to specific histone marks (H3K27ac, H3K4me1, and H3K4me3), we conducted ChIP-qPCR assays using the Magnify ChIP assay (Cat No. 492024, Thermo-Fisher, Waltham, MA, USA), following manufacturer’s guidelines. A detailed description of the method is provided elsewhere [27, 28]. Briefly, 1.5-2 × 10^6^ homozygous rs35907548 risk-“TT” (GM18603) and non-risk-“CC” (GM18624) genotype B-lymphoblastoid LCLs (Coriell) were cross-linked with 1% paraformaldehyde, washed, and sonicated. Immunoprecipitation was performed overnight at 4°C with antibodies against individual histone marks or other DNA-binding proteins pre-incubated with Dynamag magnetic A+G beads. After reverse crosslinking and elution, real-time qPCR analysis was performed with SYBR Green and primers flanking the rs35907548 region using an Applied Biosystems 7900HT qPCR machine. Statistical significance was assessed by Student’s t-test using GraphPad PRISM software, and a *p*-value < 0.05 was considered significant. Detailed methods are described in Supplementary Materials and Methods.

### 2.5 Chromatin conformation capture (3C) with quantitative PCR (3C-qPCR)

To investigate the chromatin interactions between the promoters of the target genes and the rs35907548 region in an *ex vivo* context, we utilized 3C-qPCR in LCL and Jurkat cells. Detailed protocols are provided elsewhere [27, 28]. Briefly, cells were suspended in complete media with 10% FBS (1 × 10^6^/mL media) at 70-80% confluency and cross-linked with 1% paraformaldehyde at room temperature for 10 minutes. Following quenching with 0.2 M glycine, cross-linked cells were lysed in buffer containing protease inhibitors. Cross-linked nuclei were purified, suspended in 0.5% SDS, and incubated at 62°C for 10 minutes followed by quenching with Triton X-100. Perforated nuclei were digested with 400 U *HindIII* and *DpnII* at 37°C overnight and in-nucleus ligated with T4 DNA ligase at 20°C for 4 hours. A small volume of the digested mixture was reserved to evaluate digestion efficiency. DNA was purified from ligated chromatin with proteinase K digestion, phenol-chloroform extraction, and alcohol precipitation. Purified DNA was quantified and diluted for 3C-PCR. Primers were designed to amplify several promoter regions based on restriction maps. Primers within the rs35907548 enhancer were used as common primers for other fragments. Primer sequences are in **Supplementary Table 1**. Cross-linking frequencies were calculated from PCR band intensity. Data were plotted as relative interaction frequency versus genomic distance from rs35907548.

### 2.6 CRISPR/Cas9 gene editing

To evaluate the functional consequences of the rs35907548 region, we utilized CRISPR/Cas9 gene editing. Detailed experimental protocols are available elsewhere [27, 28]. Short-guide RNA (sgRNA)/Cas9 RNP complexes were introduced into HL-60 cells by Neon Electroporation System. Genomic DNA was extracted three days post-transfection and Sanger-sequenced to verify deletion. Indel efficiency was determined using TIDE and ICE. Subsequently, pooled edited cells were cultured and harvested for gene expression measurements. sgRNA sequences are in **Supplementary Table 1**. Three experimental replicates were performed per cell type. Statistical significance was assessed by Student’s t-test using GraphPad PRISM; *p*-value of < 0.05 was considered significant.

### 2.7 Quantitative real-time PCR (qRT-PCR)

Pooled CRISPR/Cas9-edited cells were used for RNA purification using the RNA Mini-prep kit (Zymo Research). Purity and concentration were measured using a NanoDrop spectrophotometer. Approximately 700 ng total purified RNA was used to generate cDNA with the iScript cDNA Synthesis kit (Bio-Rad). cDNA was used for PCR to quantify gene expression for *SLC15A4, GLTD1*, and lncRNA AC069262.1 using qRT-PCR LightCycler 480 Instrument II (Roche) using specific primers and iTaq Universal SYBR Green Supermix (Bio-Rad). To normalize gene expression data, 18S rRNA was used as an internal control. Primer sequences are in **Supplementary Table 1**.

### 2.8 Endolysosomal pH

To assess differences in endolysosomal pH between WT and KO cells, we used the pHrodo® Red AM Intracellular pH Indicator (Thermo-Fisher) following manufacturer’s protocol. Briefly, WT and KO cells (0.2 × 10^6^ cells) were stained with 5 μM pHrodo® Red AM at 37°C for 30 minutes in 96-well plates. Cells were washed with Live Cell Imaging Media, and standard buffers containing 10 μM nigericin and 10 μM valinomycin were added to specific wells for 5 minutes to clamp intracellular pH values at 4.5, 5.5, 6.5, and 7.5. The average cellular fluorescence was measured in triplicate samples using a spectrophotometer (Synergy H1, BioTek). A standard curve was generated for WT and KO samples, showing a linear relationship between intracellular pH and relative fluorescence units.

## 3 Results

### 3.1 Systematic bioinformatics prioritizes rs35907548 as a likely regulatory SNP

We used diverse bioinformatics tools to assess the regulatory potential of all 77 high-linkage disequilibrium (LD; r^2^>80%) SNPs in and around the *SLC15A4* locus (**Supplementary Table 2**). These 77 SNPs were generated from 5 significantly associated (*p*<5 × 10^-8^) index SNPs (rs10847697/rs1385374 [29], rs12370194 [30], rs10593112 [31], and rs11059928 [32]). RegulomeDB2 ranked all SNPs for regulatory potential, prioritizing intronic rs35907548 (**Supplementary Table 2**). We used ENCODE [24]-annotated histone marks, chromatin accessibility, and RNA Pol II occupancy to identify active chromatin, within which rs35907548 lies (**Figure 1**). ENCODE annotated rs35907548 as a “distal enhancer-like” candidate *cis*-regulatory element (cCRE). We used PCHi-C chromatin-conformation data to identify several regions topologically associated with rs35907548 in multiple immune cells (**Figure 1**). To investigate the impact of rs35907548 on target gene expression, we conducted a thorough search in publicly available expression quantitative trait locus (eQTL) databases, specifically focusing on patient-derived primary immune cells [25]. Our findings indicate that rs35907548 acts as an eQTL for *SLC15A4* in various immune cell types (**Supplementary Figure 1**). Notably, plasmacytoid dendritic cells (pDCs) exhibit the most significant eQTLs on *SLC15A4*. Finally, rs35907548 is in the middle of multiple transcription factor binding sites (**Supplementary Figure 2**), where the risk allele T is universally conserved among motifs. These transcription factors include Ikaros family zinc finger protein 1 (IKZF1), IKZF3, E74-like factor 1 (ELF1), Friend leukemia integration 1 transcription factor (FLI1), and ETS1, all of which are SLE risk genes themselves [33, 34, 35, 36, 37].

**Figure 1.**
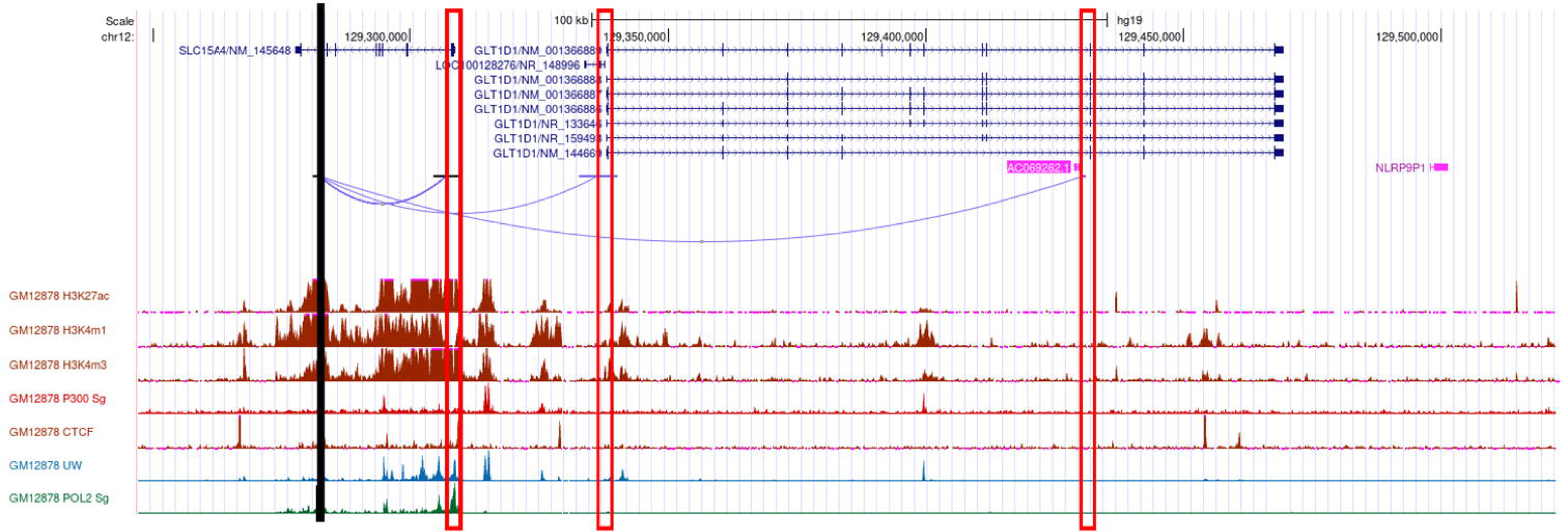
Bioinformatics. Cartoon showing the genomic location of rs35907548 (black vertical line), PCHi-C interactions from rs35907548 to the promoters (rectangle boxes) of the neighboring genes, regulatory histone marks, and Pol 2.

### 3.2 rs35907548 risk allele increases enhancer activity in diverse cell lines

We assessed allele-specific enhancer activity of rs35907548 using dual-luciferase reporter enhancer assays in both non-immune HEK293 cells and immune cells including Jurkat, HL-60, and LCL B-cells. In all cell types, the rs35907548 risk allele (TT) exhibited notably higher enhancer activity compared to the non-risk allele (CC) (p-values: < 0.0001, < 0.001, 0.016, 0.0016; as shown in **Figure 2a**). As negative controls, we selected two non-coding SNPs, rs12831705 and rs34616325. Both SNPs, at distances of 12.7 kb and 25.5 kb from the target SNP, respectively, are scored “1f” (i.e., predicted to have a similar regulatory potential as rs35907548; **Supplementary Table 2**) by RegulomeDB and are in strong linkage disequilibrium (LD) with rs35907548. Our results show insignificant regulatory activity for both negative control SNPs in HEK293 and LCL (**Supplementary Figure 3**), underscoring the effects

**Figure 2.**
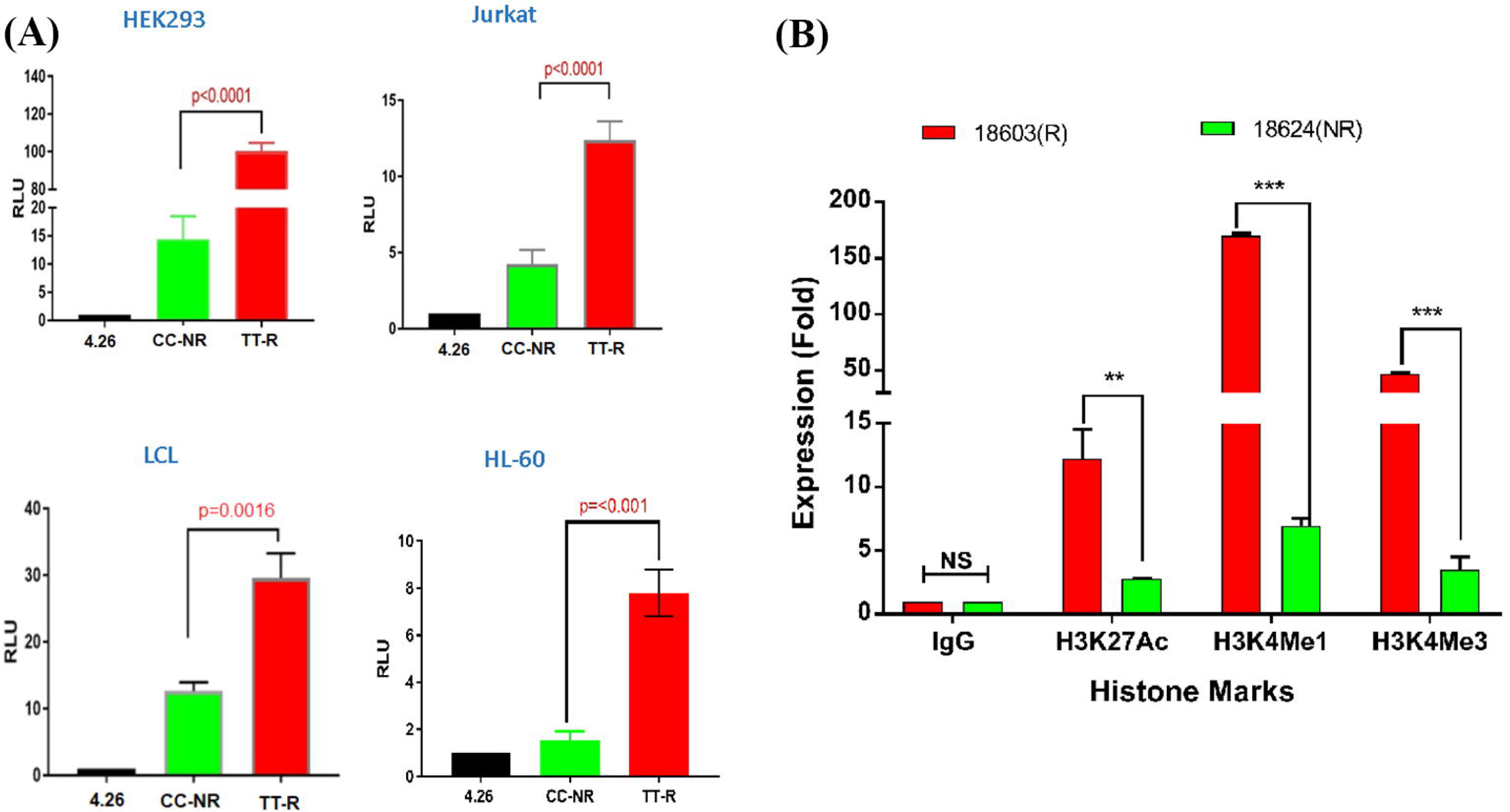
Allelic-specific regulatory effects of the rs35907548enhancer. **(A)** Luciferase reporter assays on 4 cell types. **(B)** ChIP-qPCR with risk (R) and non-risk (NR) LCLs.

Using ChIP-grade antibodies, we analyzed the specific binding patterns of three regulatory histone marks—namely, H3K27ac, H3K4me1, and H3K4me3—at the SNP site within its biological context. The quantification of binding was conducted through ChIP-qPCR (**Figure 2b**). Across all three marks, we observed a distinct and heightened level of binding in GM18603 (the risk TT genotype) when compared to GM18624 (the non-risk CC genotype) (H3K27ac: p = 0.002, ∼4-fold increase; H3K4me1: p < 0.001, ∼20-fold increase; H3K4me3: p < 0.001, ∼13-fold increase). This substantiates the prevailing notion that the SNP is situated within an active enhancer region, where intricate interactions involving RNA polymerase, histone marks, chromatin modulators, and allele-specific components collectively oversee the orchestration of transcriptional control at these specific loci.

### 3.3 The rs35907548 enhancer establishes long-range chromatin interactions with target gene promoters

Given that rs35907548 demonstrated allele-specific enhancer activity across all examined cell lines, we set out to explore its potential involvement in establishing promoter-enhancer connections through chromatin interactions. To explore these interactions, we performed 3C-qPCR experiments between the SNP and gene promoters (**Figure 2b**). Our findings aligned with previous eQTL and pCHiC data, unveiling interactions between the enhancer-SNP region and the promoters of *SLC15A4, GLT1D1*, and LncRNA *AC069262.1* (as illustrated in **Figure 1**). These collective results imply that the enhancer region encompassing rs35907548 could play an active regulatory role in governing the expression of *SLC15A4, GLT1D1*, and LncRNA *AC069262.1*.

### 3.4 Validating transcriptional effects with CRISPR-based genome editing

To validate the transcriptional regulatory effects of the rs35907548 locus, we used three short-guide RNAs (sgRNAs) to delete ∼140 bases around rs35907548 in HL-60 cells (**Figure 4a** and **Supplementary Figure 4**). The deletion was confirmed with Sanger sequencing, and ICE analysis demonstrated high indel efficiency (63%) of pooled cells (data not shown). Subsequently, we determined expression levels of *SLC15A4, GLT1D1*, and lncRNA *AC069262*.1 in KO and WT cells. Expression levels were lower in KO cells than WT (∼45%, *p* < 0.0001; ∼25%, *p* < 0.01; and ∼35%, *p* < 0.001, respectively; **Figure 4b**). These findings provide further evidence that rs35907548 exerts regulatory effects to influence the expression of *SLC15A4* and other target genes through its enhancer activity.

**Figure 3.**
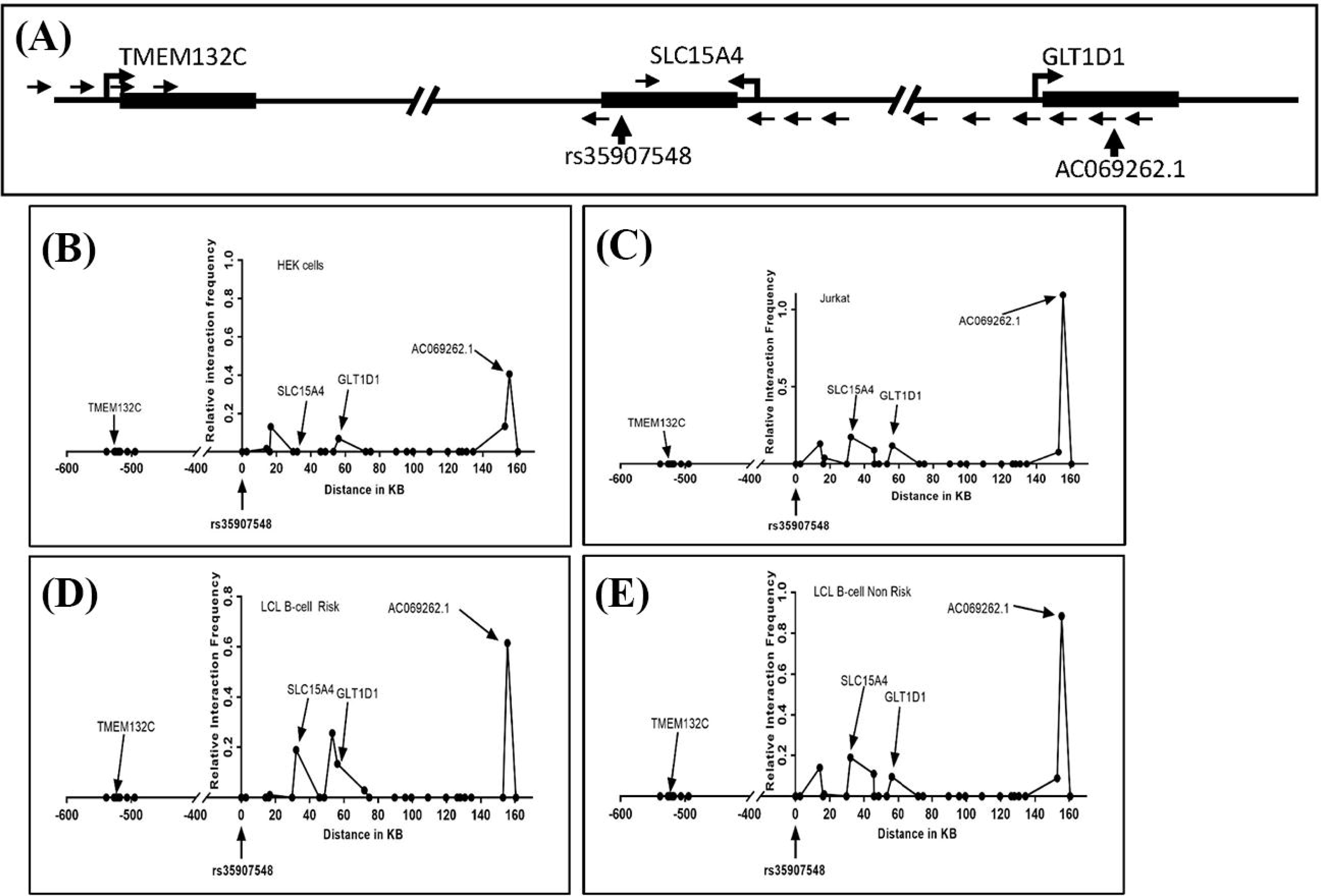
3C interaction analysis. **(A)** Schematic of the rs35907548 region and neighboring genes. SacI restriction enzyme sites were used to design primers. Small arrows represent primer locations and orientations. The common primer at the rs35907548 region is common for all other primers at *SLC15A4*, *GLT1D1* and *TMEM132C* promoter regions. Big arrowheads represent transcriptional start sites (TSSs). **(B-E)** Graphs show relative interactions of rs35907548 regions with different genomic regions in HEK (**B**), Jurkat **(C)** and EBV-transformed LCL B-cells **(D, E)**. The relative interaction frequency for each set of primers represents the intensity of PCR. X-axis shows genomic distance (in kb) in forward and reverse directions from rs35907548 (0 kb).

**Figure 4.**
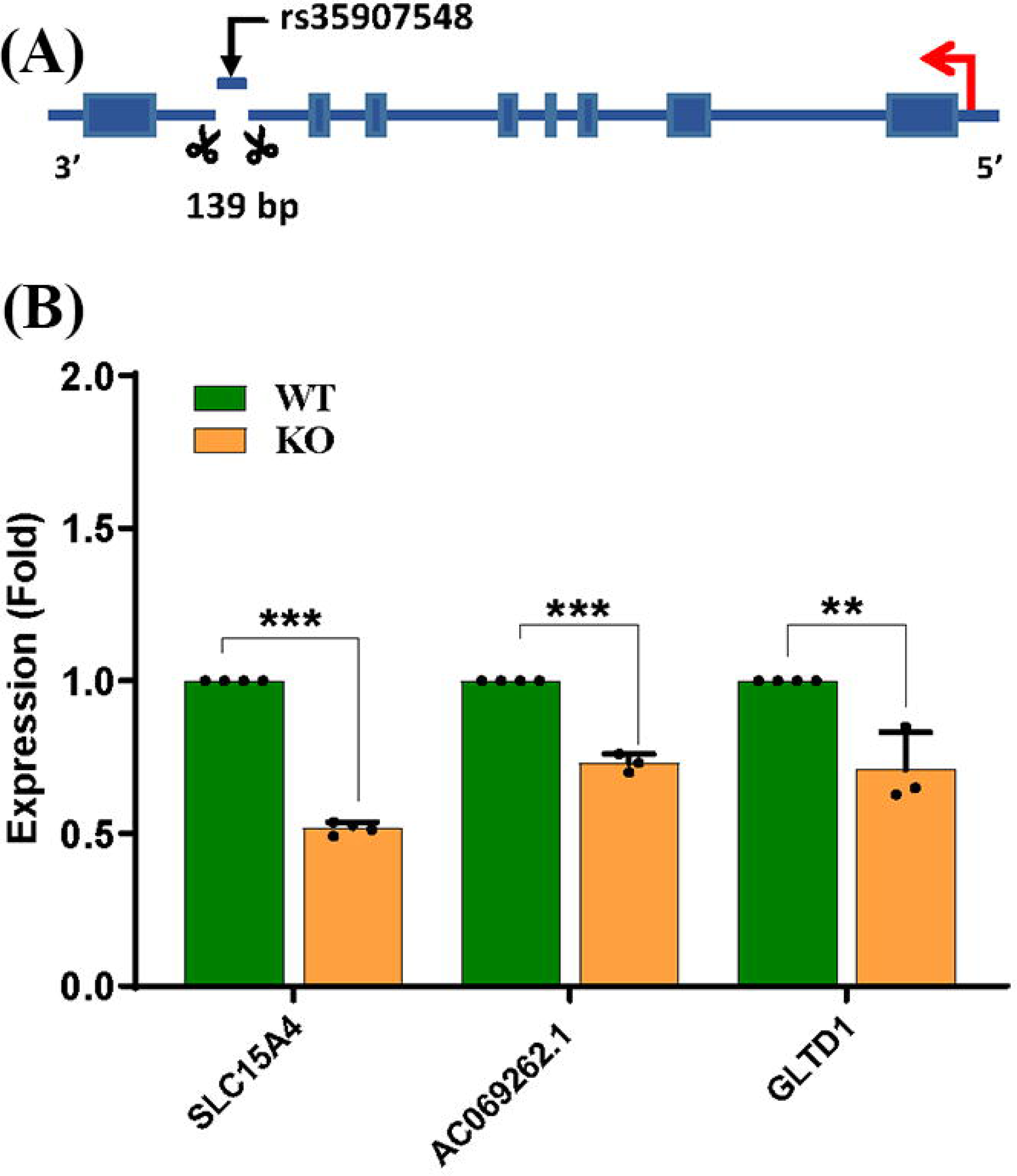
CRISPR/Cas9-based enhancer deletion. **(A)** CRISPR-based deletion, **(B)** qPCR-based target gene expression between WT and KO.

### 3.5 Impact of the rs35907548 enhancer on endolysosomal acidification

*SLC15A4* plays a crucial role in maintaining the acidic environment of the endolysosomal compartment [38]. We found that KO cells expressed less SLC15A4; consistent with this, KO cells showed a dramatic increase over WT in endolysosomal pH (5.3 *vs*. 4.5, *p* < 0.01; **Figure 5**). These data underscore the crucial link between SLC15A4, maintenance of the acidic environment in endolysosomal compartments, and subsequent effects on cellular pH regulation.

**Figure 5.**
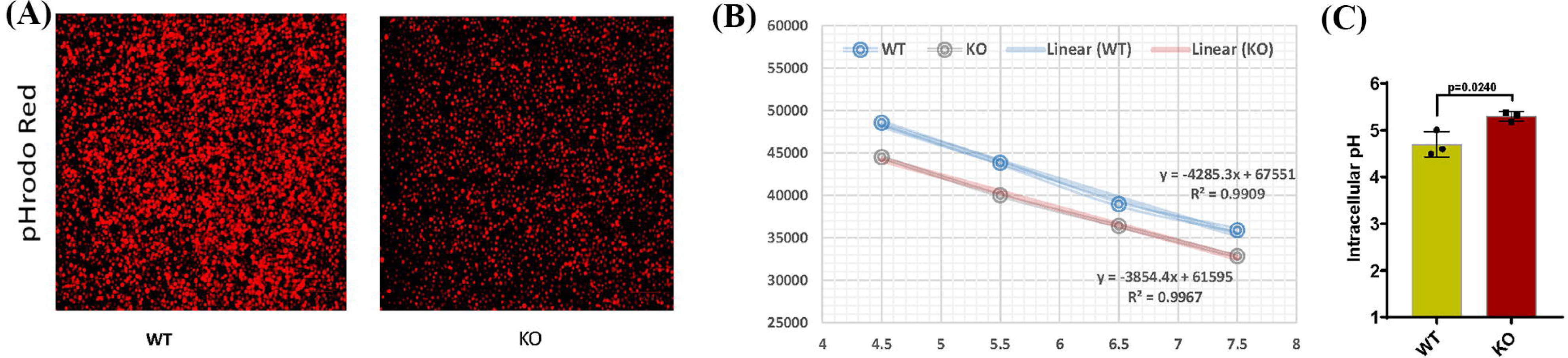
Comparing the endolysosomal acidification between WT and KO cells. **(A)** pHrodo™ Red AM-stained WT and KO cells under a fluorescent microscope. **(B)** Standard curve for WT and KO cells using pHrodo™ Red AM with Intracellular pH Calibration Buffer Kit for the translation of fluorescence ratios into pH. **(C)** pH determined for WT and KO using standard curve and manufacturer’s protocols. RFU = Relative Fluorescence Units.

## 4 Discussion

GWAS analyses invariably produce peaks of association; these peaks are often quite large, covering many genes and 1000s of SNPs. Sifting through these genes and SNPs to identify the true causal variants and their mechanisms of action from broad GWAS peaks is challenging [39], but an essential prerequisite for understanding disease risk and opportunities for diagnosis and treatment. Several studies have linked *SLC15A4* SNPs to SLE risk in Asians and Europeans [18, 19, 40], albeit with no experimental testing.

Here, we localize *SLC15A4* SLE risk to the intronic SNP rs35907548, at the center of an enhancer modulating expression of – and physically interacting with – *SLC15A4* and nearby genes. Dual-luciferase reporter assays revealed strong enhancer activity, particularly of the risk “TT” allele, in diverse cell types. The risk allele consistently showed substantially higher binding to active histone marks. We confirmed the activity of the rs35907548 enhancer in cells by creating CRISPR/Cas9 knock-out cells, which showed significantly lower levels of *SLC15A4*, *GLT1D1*, and the uncharacterized lncRNA *AC069262.1.* SLC15A4 is a histidine transporter located primarily on the endolysosomal membrane, where it plays several pivotal roles including maintenance of endolysosomal acidification [19, 38], thus regulating protein degradation, inflammation, endocytosis, and autophagy, among other processes [41, 42, 43]. Accordingly, KO cells failed to properly acidify the endolysosome, demonstrating the profound effects of the rs35907548 enhancer on cell physiology.

Several studies have suggested SLC15A4 as a potential therapeutic target for systemic lupus erythematosus (SLE) and other autoimmune diseases [21, 38, 44]. In mouse models, the absence of *SLC15A4* has been shown to confer resistance to the development of multiple autoimmune diseases, including dextran sulfate sodium (DSS)-induced colitis (wherein *Slc15a4*^-/-^ pDCs fail to produce IFNα upon TLR7 agonist R848 stimulation) and the Faslpr model of SLE [18]. This observation is significant as TLR7/8 inhibitors are currently in clinical trials for SLE [19]. These findings collectively underscore the dysregulation of lysosomal pH, particularly deacidification, in autoimmune diseases such as SLE. This dysregulation is closely linked to the activity of lysosomal hydrolases.

Interestingly, two widely used anti-malarial drugs, hydroxychloroquine and chloroquine, are commonly used to treat SLE. These drugs have many cellular targets and complicated mechanisms of action. The weak base drugs accumulate in acidic compartments like the lysosome, where they increase pH by becoming protonated. The drugs also directly inhibit lysosomal hydrolases, increasing pH by a further 2 units. Finally, they block stimulation of Toll-like receptor 9 (TLR9) family members, decreasing innate immune reactivity. Together, these effects decrease lysosome activity and inflammasome activation and mitigate inflammation. The precise mechanisms through which these drugs benefit SLE patients are still being investigated, and other cellular targets may be discovered. Despite this incomplete understanding, these drugs are critical tools in managing SLE and related inflammatory diseases [45, 46, 47].

Crucially, our study identifies *SLC15A4* as a genetic locus linked to substantially elevated SLE risk. Given its central role in governing endolysosomal pH and activity, this genetic variation likely contributes to SLE susceptibility, aligning well with the fact that endolysosomal modulator drugs are essentially the only SLE drugs of note. Our findings, as depicted in Figures 2a and 4, point to heightened SLC15A4 expression in SLE patients. Importantly, a recent study [48] lends additional support to this assertion. Despite its reliance on a limited sample size, the study reports significantly elevated *SLC15A4* mRNA in PBMCs from SLE patients compared to healthy controls. This observation bolsters the association between *SLC15A4* expression and autoimmune diseases such as SLE [49].

rs35907548 lies in the middle of multiple conserved transcription factor binding sites, including those for SLE risk genes and white blood cell factors IKZF1, IKZF3, ELF1, FLI1, and ETS1; the ancestral risk allele T is universally present in the binding motifs. It is likely that proper binding of these transcription factors is critical to enhancer function, and indeed the risk allele T shows much higher levels of active chromatin marks and enhancer activity, implying a heightened regulatory role.

The convergence of various elements – including the existence of evolutionarily conserved binding sites, the participation of transcription factors associated with immune processes and SLE, and the distinct presence of the risk allele within these motifs – collectively implies a complex interplay that likely underlies the regulatory function of the enhancer. These findings emphasize the intricate and multifaceted nature of genetic aspects of disease susceptibility and cellular function.

The adjacent locus Glycosyltransferase 1 Domain Containing 1 (*GLT1D1*) has also been flagged as an SLE risk locus [50]. GLT1D1 targets programmed cell death-ligand 1 (PD-L1) for glycosylation [51]; glycosylated PD-L1 is strongly immunosuppressive and correlates with B-cell non-Hodgkin’s lymphoma progression. In addition to the links to endolysosomal acidification, TLR signaling, and autophagy through SLC15A4, future experiments will investigate the contribution of rs35907548 and other SNPs to SLE risk through *GLT1D1*. The role of lncRNA *AC069262.1* in biological processes and diseases, including SLE, remains largely unexplored. Further investigations are required to uncover its potential significance and contribution to SLE.

In the present study, our primary objective was to define susceptibility regions likely to contain functional SNP(s) linked to increased lupus risk. Leveraging bioinformatics and experiment, we identified rs35907548 within an active enhancer with the capacity to modulate expression of target genes with critical immune functions. Nevertheless, we acknowledge a limitation of our study, namely that we have not explicitly demonstrated effects of the single base-pair change on target gene expression, endolysosomal pH, or other cellular properties. To address this limitation, advanced techniques like CRISPR base-editing or prime-editing will be required. We have demonstrated numerous specific effects of the single base-pair change (transcription factor binding, enhancer activity, chromatin contacts) from both in vitro experiments and from data acquired from *in vivo* samples, which are strongly consistent with the cellular effects that we observe from CRISPR deletion of the rs35907548 locus. Future studies will validate the cellular effects of the single base-pair change, allowing stronger conclusions to be made about the effect of the rs35907548 SNP on SLE development and progression.

In summary, our study both identifies a SNP causally underlying SLE risk association at the *SLC15A4* locus and establishes an analytical and experimental framework for studying risk SNPs at *SLC15A4* and other loci. The results and framework may contribute to future investigation of therapeutic interventions and diagnostics.

## 5 Data availability statement

All supporting data is available upon request from the corresponding author.

## 7 Author Contributions

SKN conceived and designed the study. MKS, GPM, HRR conducted and validated the experiments. MFN, LLL, SKN organized the database and conducted bioinformatics analyses. SKN, GPM, MKS, LLL wrote the manuscript. All authors contributed to manuscript revision, and read and approved the submitted version.

## 9 Acknowledgments

The authors wish to sincerely express their appreciation to Dr. Valerie Harris for her invaluable advice on setting up the pHrodo experiments. Furthermore, they extend their thanks to Ms. Louise Williamson for her technical assistance during the preparation of the manuscript.

## 10 Figure Legends

**Supplementary Figure 1.**
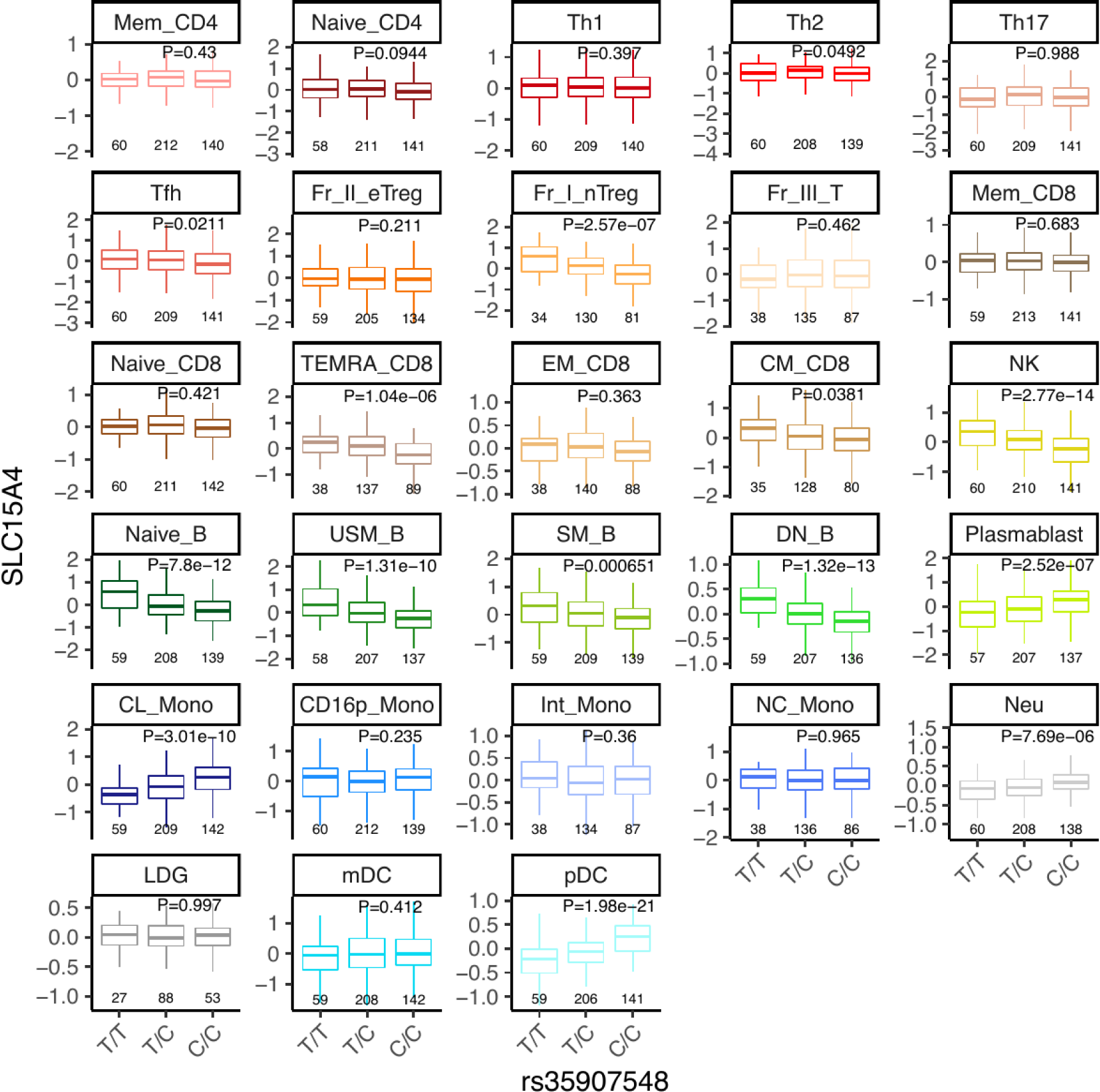
Immune cell type-specific eQTLs for rs35907548. The genotypes and their corresponding sample sizes are shown below each box plot. (T/T) is the risk allele for rs35907548. USM = unstimulated, SM = stimulated, DN = double negative.

**Supplementary Figure 2.**
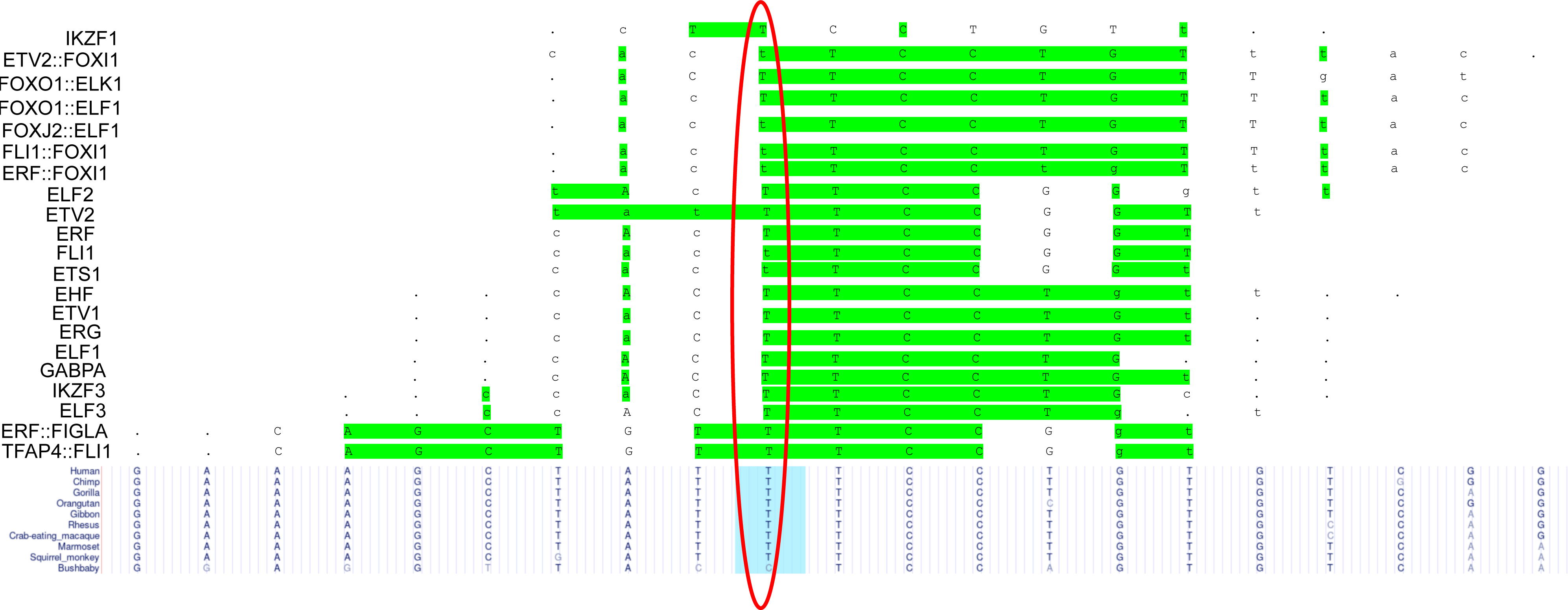
Transcription factor binding sites overlapping rs35907548. The ancestral risk allele T (red oval) produces more favorable motifs for all TFs. Green highlights bases where the consensus motif is present.

**Supplementary Figure 3.**
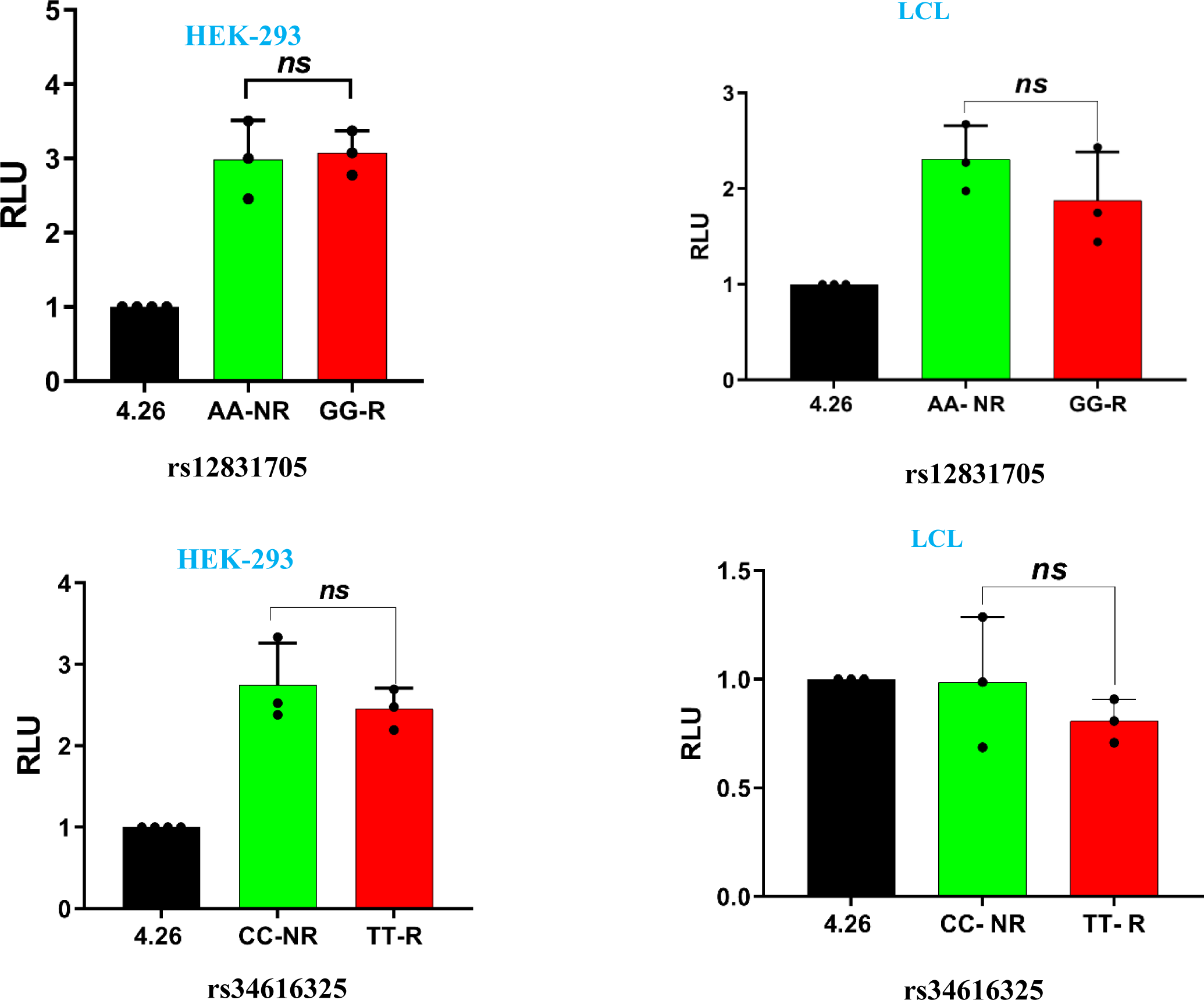
Negative control for assessing allelic-specific enhancer activity. Luciferase reporter assays were conducted on sequences surrounding the predicted inactive SNPs rs12831705 and rs34616325 in HEK-293 and LCL cells, revealing non-significant allele-specific regulatory effects.

**Supplementary Figure 4.**
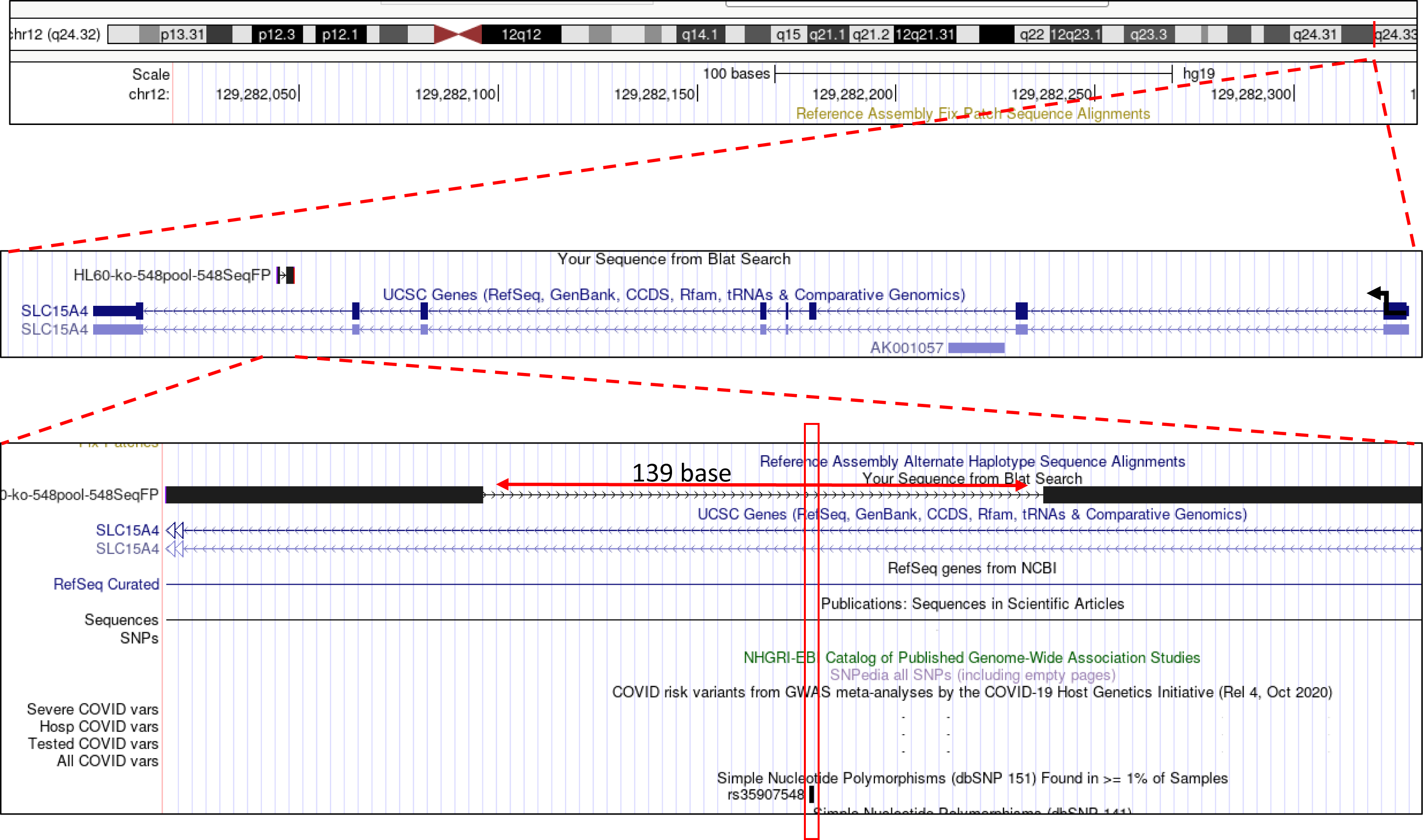
CRISPR/Cas9-based enhancer deletion of 139 bp region containing rs35907548.

